# Differential contributions of human oligosaccharyltransferase complexes OST-A and OST-B to HIV-1 envelope glycoprotein glycosylation

**DOI:** 10.1101/2025.09.03.674041

**Authors:** Tugba Atabey, Ronald Derking, Maddy Newby, Joey Bouhuijs, Jonne Snitselaar, Yoann Aldon, Joel D. Allen, Max Crispin, Rogier W. Sanders

## Abstract

N-linked glycosylation of glycoproteins during synthesis in the endoplasmic reticulum (ER) is mediated by oligosaccharyltransferase (OST) complexes OST-A and OST-B that have different catalytic subunits STT3A and STT3B, respectively. OST-A acts cotranslationally, while OST-B adds glycans posttranslationally. While there is redundancy between these two enzymes, it is unclear how they both contribute to glycosylation of the densely glycosylated HIV-1 envelope glycoprotein complex (Env). We found that knocking out STT3A had a profound negative impact on HIV-1 virus production and infectivity while STT3B ablation had no such effect suggesting that STT3A is more important than STT3B for Env glycosylation and preserved function. STT3A/3B knockout (KO) affected the neutralization sensitivity to broadly neutralizing antibodies (bNAbs) in a strain-specific manner with STT3A-KO increasing susceptibility to VRC01 bNAb for the tested HIV-1 strains. In contrast, for the BG505 strain A virus, it conferred increased resistance to glycan-dependent bNAbs 2G12 and PGT128. For other HIV-1 strains, STT3B-KO also led to resistance to glycan-dependent bNAb PGT151. Site-specific glycan analysis of recombinant Env proteins revealed that STT3A-KO reduced glycan occupancy of potential N-linked glycosylation sites (PNGS) more globally than STT3B-KO, with certain acceptor sites, including N234 and N386, showing STT3A dependence. In contrast, STT3B-KO appeared to have a more pronounced effect on gp41 glycosylation, suggesting that PNGS located near the C-terminus are more dependent on STT3B. Defining the roles of the OST-A and OST-B complexes in HIV-1 Env glycosylation may bring critical information for the development of methods to control PNGS glycan occupancy of recombinant glycoprotein immunogens.

## INTRODUCTION

N-linked glycosylation of polypeptides is crucial for proper folding, stability, protein trafficking and secretion (1–3). HIV-1 envelope glycoprotein (Env) is among the most heavily glycosylated proteins found in nature, with glycosylation playing a critical role in viral infectivity, immune evasion, and host interactions (4–8). N-linked glycans are attached to potential N-linked glycosylation sites (PNGS) on each protomer, totaling to up-to 100 PNGS per Env trimer and accounting for approximately half the molecular mass of the external domains of Env. HIV-1 Env gp120 subunits are densely glycosylated with up to 35 PNGS per gp120 protomer, while the gp41 subunits typically harbour 4 PNGS per protomer (7, 8).

PNGS constitute of asparagine (N) residues in NxT/Sx sequons, where x is any amino acid except proline. The presence of proline at the x position disrupts this sequon and prevents glycosylation, while residues at the second (x), third (T/S), as well as those flanking the sequon, influence the efficiency of N-linked glycan addition (9, 10). Oligosaccharyltransferase (OST) complexes add glycan precursor molecules (GlcNac_2_-Man_9_-Glc_3_; where GlcNAc, Man, and Glc, are N-acetylglucosamine, mannose, and glucose, respectively) from a dolichol-pyrophosphate carrier to the NxT/Sx motifs on newly synthesized proteins in the lumen of the endoplasmic reticulum (ER) (11, 12). OST complexes are heterooligomeric transmembrane enzyme complexes embedded in the ER membrane. In human cells, two different OST complexes are expressed and located in this compartment: OST-A and OST-B. Both OST-A and OST-B complexes regulate cotranslational and posttranslational glycosylation, respectively (13). OST-A and OST-B contain different catalytic subunits referred to as STT3A and STT3B, respectively, and share a set of non-catalytic subunits –including ribophorin I (Rb1), ribophorin II (Rb2), OST48, DAD1, and OST4– plus complex-specific subunits that are part of the OST complex (i.e. DC2 for OST-A and MagT1 for OST-B) (13–15). STT3A and STT3B enzymes exhibit preferences for certain PNGS motifs. As a consequence, OST isoforms preferentially recognize certain NxT/Sx motifs based on the amino acid context adjacent to PNGS targeted motifs. Additionally, steric hindrance or conformational shielding of N-glycosylation sites can limit the ability of OST isoforms to access and glycosylate certain PNGS (16, 17).

The trimeric HIV-1 Env glycoprotein, which is the sole target for broadly neutralizing antibodies (bNAbs) that arise during the natural infection (18, 19), has become the major focus for HIV-1 vaccine development research (20–22). Env, the sole surface protein on HIV-1 virions, is the only virion-associated protein modified by glycosylation; therefore, disrupting glycan addition is unlikely to affect any other viral components (23, 24). Functional Env trimers are derived from gp160 polypeptide precursor chains, which are cleaved into gp120 and gp41 subunits by host proteases (i.e. furin), associated into type I fusion glycoprotein timers and mediate viral entry into host cells (25, 26). The N-linked glycans that decorate the Env trimer play crucial roles in the viral life cycle, such as binding to lectin receptors, Env protein folding, trafficking between cellular compartments, and immune escape by shielding underlying conserved protein epitopes (7, 19, 27, 28). The glycan addition and composition of recombinant HIV-1 Env trimers can be different from their viral counterparts. PNGS occupancy at some specific PNGS is generally lower on recombinant Env trimers (8, 29, 30). The absence of glycans at these sites can result in holes in the glycan shield, ’glycan holes’, forming neo-epitopes that may play a significant role in shaping the antibody response against Env. Glycan holes can form due to the absence of conserved PNGS. For example, in the BG505 virus and its corresponding recombinant Env trimers, the conserved glycosylation sequons at positions N241 and N289 are not present which creates an immunodominant glycan hole (31, 32). Moreover, glycan holes can appear through incomplete glycosylation of existing functional PNGS motifs when OST-A and OST-B fail to glycosylate a given PNGS (13, 33, 34). It has been shown that PNGS underoccupancy in hypervariable V1 and V2 loops of soluble BG505 SOSIP.664 trimers, found at the trimer apex, can create artificial glycan holes in some Env protomers leading uneven underoccupancy of specific sites per trimer protomers (8). Similarly, when PNGS at position N611 is underoccupied on BG505 SOSIP soluble immunogens, the resulting antibody responses elicited are able to neutralize viruses lacking a glycan at this position but do not display neutralization activity against wild type (WT) viruses. Furthermore, nsEMPEM analysis of the sera from BG505 SOSIP vaccinated rhesus macaques revealed the immunodominance of two epitope clusters, the V1/V2/V3 region and the trimer base, of which in particular the latter is underglycosylated (31, 32, 35, 36). These glycan holes may distract from more desirable responses and impede the development of neutralization breadth (31, 32, 37). Understanding the limiting factors behind glycan holes on HIV-1 Env recombinant trimers may provide valuable insights for improving immunogen design and vaccine development.

Recently, the CRISPR/Cas9 gene-editing system was used to generate HEK293-derived knockout (KO) cell lines that are deficient for a single catalytic subunit, STT3A or STT3B (38). In the present study, these KO cell lines were used to study the contribution on STT3A or STT3B to PNGS occupancy on viral and recombinant Env glycoproteins. We studied the impact of STT3A/STT3B-KO on viral infectivity of diverse HIV-1 strains. The recombinant Env glycoproteins were produced in these cell lines for site-specific glycan analysis to interrogate possible STT3A or STT3B-dependent PNGS. Our results therefore contribute to increasing fundamental understanding of viral glycoprotein dependency on OST and highlights potential paths to explore in order to develop methods to better control glycan occupancy on Env-based and other type 1 fusion immunogens.

## RESULTS

### Knocking out STT3A greatly reduces HIV-1 infectivity

To explore the roles of the two OST complexes in HIV-1 biology, intact virus from the well-studied BG505 strain was produced in WT, STT3A- and STT3B-KO HEK293T cells as previously described (20)(Figure 1A&B). A p24 antigen capture ELISA revealed that virus production was severely impaired in STT3A-KO cells (Figure 2A). This is consistent with previous observations that HIV-1 Env biosynthesis problems can disrupt overall virus production and egress from infected or producer cells (39–42). We then assessed virus infectivity of BG505 strain by measuring its TCID_50_ using the TZM-bl reporter assay, where this modified cell line can be infected by HIV-1 thanks to transgenic expression of CD4, CCR5 and CXCR4 receptors and co-receptors necessary for Env mediated viral entry. Impairment of the OST-A activity by knocking out STT3A caused a dramatic reduction of HIV-1 infectivity *in vitro*. However, the absence of OST-B activity did not have an appreciable impact, suggesting that N-linked glycosylation of HIV-1 Env native protein is more dependent on OST-A than on OST-B (Figure 2B).

**Figure 1.**
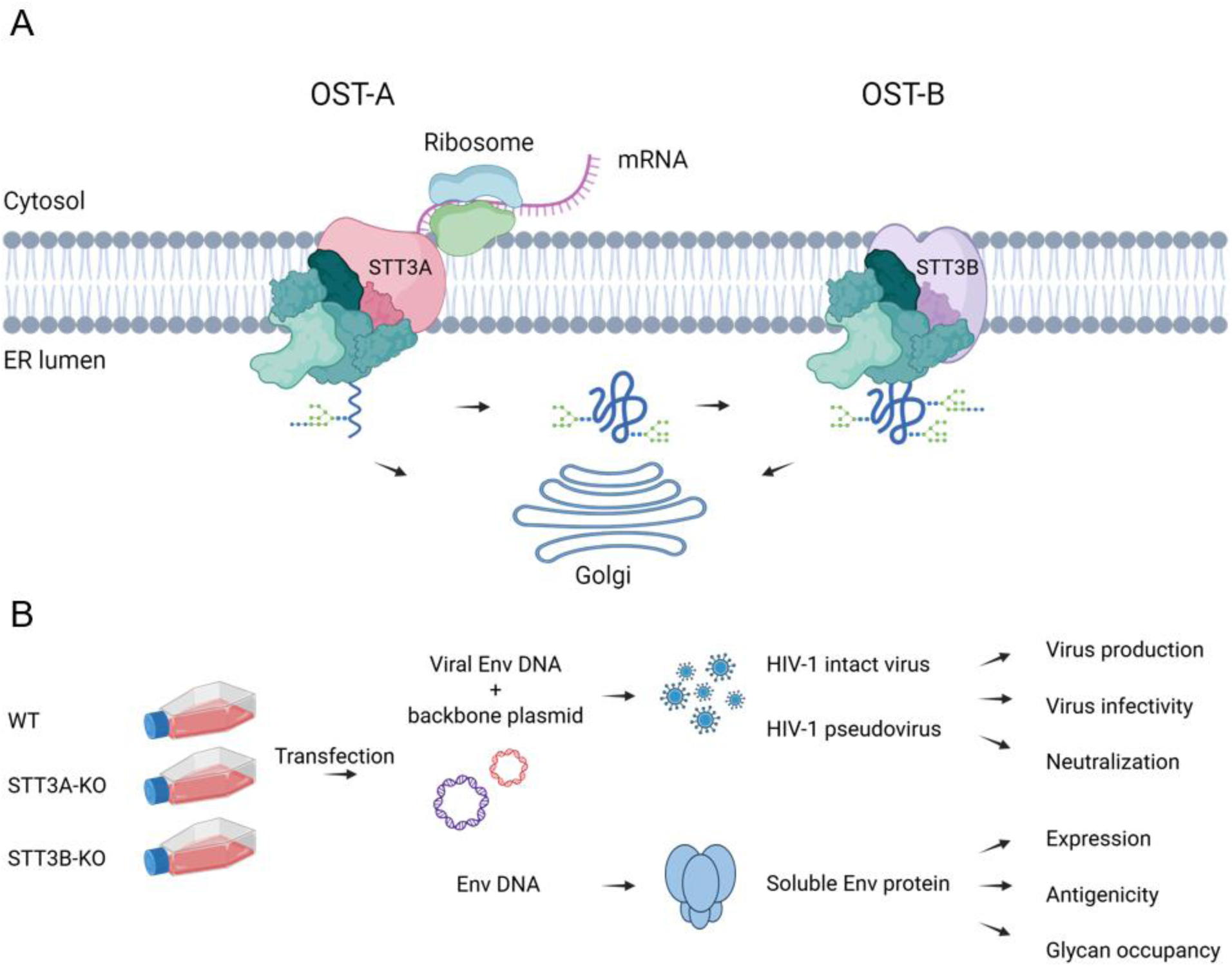
Contribution of OST-A and OST-B complexes to HIV-1 Env glycosylation. **(A)** Schematic representation of OST-A/OST-B-mediated glycosylation in the endoplasmic reticulum (ER). OST-A predominantly catalyzes N-linked glycosylation cotranslationally, adding glycans as the polypeptide is synthesized. In contrast, OST-B acts posttranslationally to glycosylate sites that have been skipped by OST-A, ensuring more complete glycan occupancy. Following initial glycosylation in the ER, polypeptides are trafficked through the Golgi apparatus, where further glycan maturation and processing occur. **(B)** Schematic representation of the experimental setup where WT, STT3A-KO, and STT3B-KO HEK293T cells were used to produce HIV-1 virus and recombinant soluble Env trimers. Figures were created using the commercial scientific illustration service BioRender.

**Figure 2.**
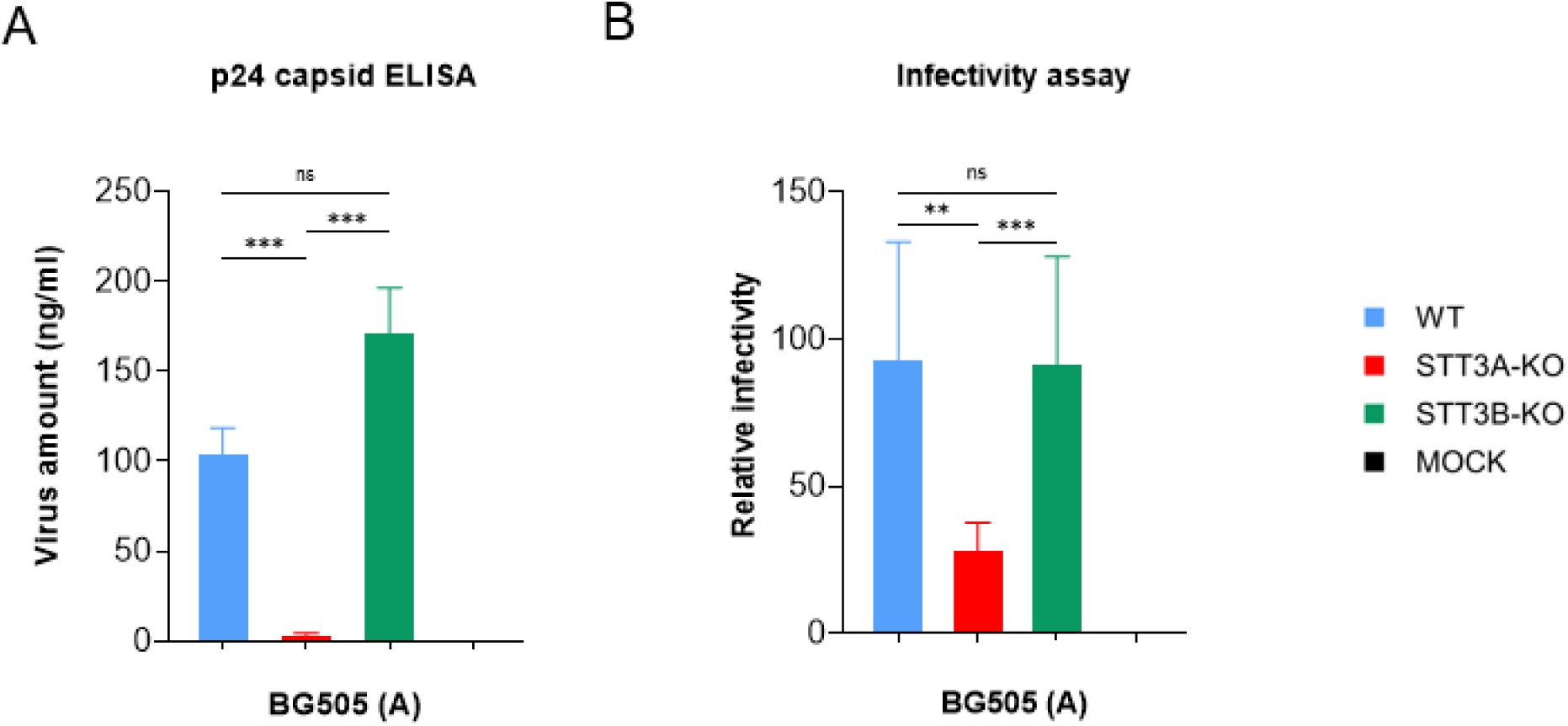
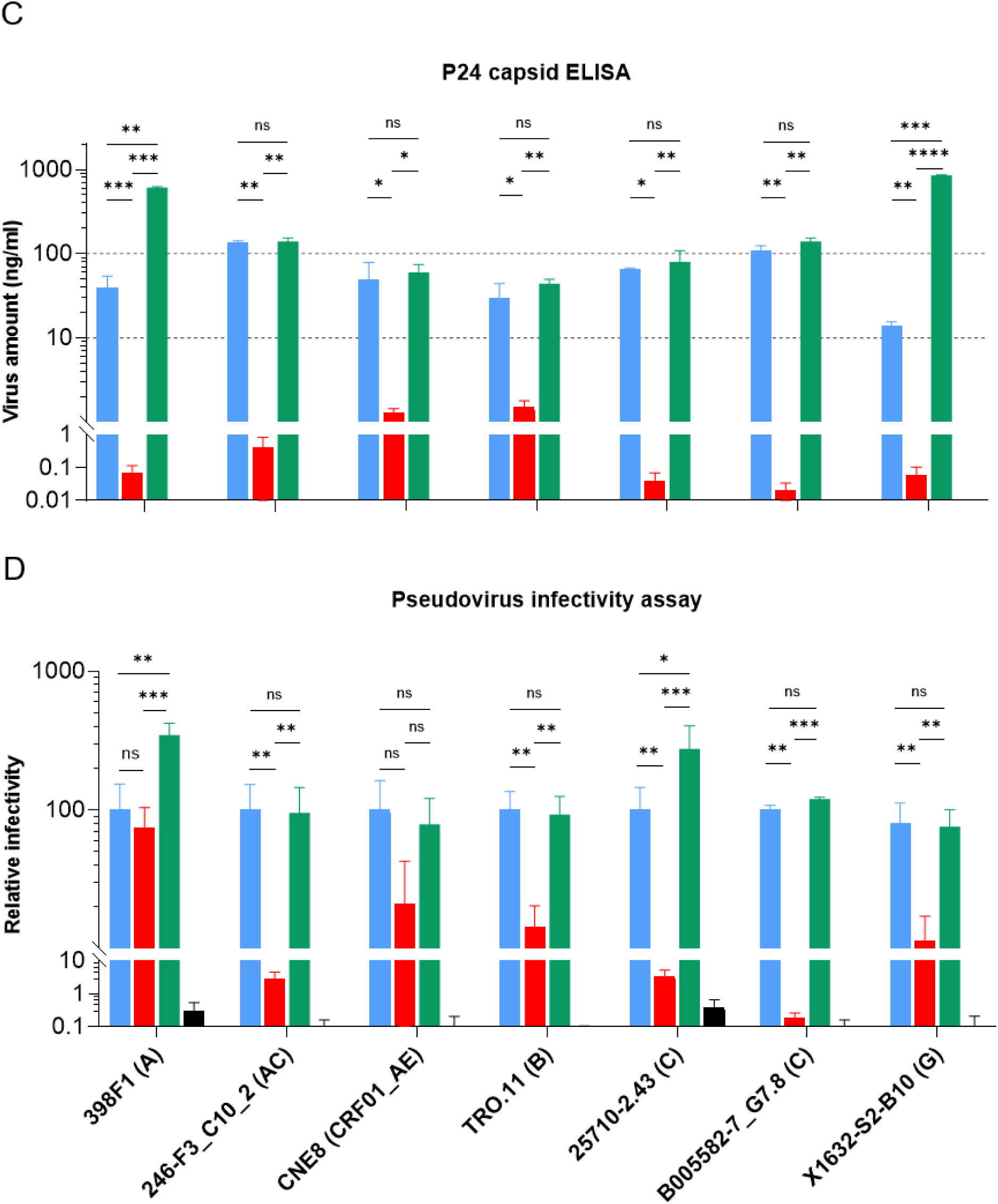
Effect of STT3A and STT3B knockouts on in vitro HIV-1 viral production and infectivity. **(A)** Quantification of p24 capsid of BG505 viruses produced using a molecular clone in three different cell lines (blue: wild type (WT), red: STT3A-KO, green: STT3B-KO). The viral productions were quantified by measuring p24 capsid concentrations using a p24 standardized ELISA. **(B)** Relative infectivity of BG505 virus produced in STT3A-KO and STT3B-KO cell lines. Infectivity was assessed using TZM-bl reporter cells, luciferase activity measured and infectivity values normalized to viruses produced in WT cells. **(C)** Quantification of p24 levels by ELISA for pseudotyped HIV-1 viruses derived from different clades/strains and produced in various cell lines. **(D)** Relative infectivity of pseudoytyped HIV-1 viruses. The measured luciferase activity was normalized to that of the WT virus. For HIV-1 strains, the clade they belong to is indicated in parenthesis. Bars represent the mean ± standard deviation from at least three independent experiments. Statistical significance between conditions was assessed using an unpaired t-test. Significance levels are indicated as follows:*, P < 0.05; **, P < 0.01; ***, P < 0.001; ****, P < 0.0001.

Besides BG505, other HIV-1 strains derived from clades A, B, C, G and recombinant forms AC and AE were also produced in these cell lines and assessed for virus production and infectivity. While the above studies on the BG505 strain were performed with intact virus, the following studies were performed with pseudotyped viruses (see materials and methods for details). Our results, using pseudotyped viruses, demonstrated that STT3A-KO significantly impaired viral production for all strains tested. In contrast, ablation of STT3B showed no deleterious effect on production and infectivity for all pseudoviruses tested (Figure 2C-D). Unexpectedly, we observed that for 398F1 (clade A) and X1632-S2-B10 (clade G) strains produced in STT3B-KO pseudoviruses production was significantly increased (Figure 2C). When studying the effect of STT3A or STT3B-KO on infectivity of these diverse HIV-1 strains, it appeared that knocking out of STT3A significantly reduced infectivity for most strains, with the exception of the 398F1 (clade A) and CNE8 (clade CRF01_AE) pseudoviruses, although we noted a trend towards reduction in infectivity for CNE8 (Figure 2D). Impairing STT3B had no negative impact on infectivity of any strain. In fact, the infectivity of the 398F1 and 25710-2-43 strains was enhanced when produced in STT3B-KO cells. The increased viral infectivity observed for the 398F1 virus is likely to be a consequence of the enhanced production observed for this pseudovirus. However, the enhanced production of the X1632-S2-B10 strain did not result in enhanced infectivity, while 25710-2- 43 infectivity was enhanced when produced, STT3B-KO cell, while virus production was not affected. These observations suggest subtle strain-specific differences in glycosylation by OST-A and OST-B, but overall we conclude that OST-A is critical for infectivity of most virus isolates, while OST-B enzymatic complex play a more subtle role in modulating viral yield and infectivity.

### Knocking out STT3A affects HIV-1 sensitivity to bNAbs

Next, we assessed whether manipulating OST-A and OST-B activity influenced sensitivity to bNAbs, in particular ones that are heavily dependent on specific Env PNGS glycosylation status. The following bNAbs were selected: PGT145 binding a quaternary epitope at the trimer apex and requiring glycans at N156 and N160; PGT151, targeting a quaternary epitope at the gp120/gp41 interface involving glycans at N611 and N637; PGT128 against the V3-N332 epitope cluster dependent on N295, N301 and N332; 2G12 targeting an epitope that consists exclusively of glycan and involves glycans at N295, N332, N339 and N386; and finally VRC01 directed against the CD4-binding site (CD4bs) and not requiring glycans for binding (43–47). In our assays, we observed that neutralization sensitivity of BG505 virus to some bNAbs was substantially altered upon manipulation of OST-A (Figure 3A). Most notably, STT3A-KO cell line production rendered the BG505 virus more susceptible to VRC01 neutralization. This result could be explained by the STT3A defect resulting in the absence of one or more glycans surrounding the CD4bs, allowing better access to the VRC01 epitope. Indeed, the site-specific occupancy data described below support that hypothesis. Another notable finding consisted in that the BG505 virus produced in STT3A-KO cells was more resistant to neutralization by the glycan dependent bNAbs 2G12 and PGT128, suggesting that one or more of the glycans involved in those targeted epitopes was/were not attached to the protein. The site-specific occupancy data (see below) indicated that the N339 and N386 PNGS were underoccupied as a consequence of STT3A impairment. With regard to quaternary- and glycan-dependent PGT151 and PGT145 bNAbs, no detectable difference in neutralization sensitivity was observed for BG505 virus. We noted that the ablation of STT3B activity had no noticeable effect on BG505 neutralization sensitivity, further emphasizing the critical role of STT3A compared to STT3B in linking glycans to Env glycoprotein in the viral membrane context.

**Figure 3.**
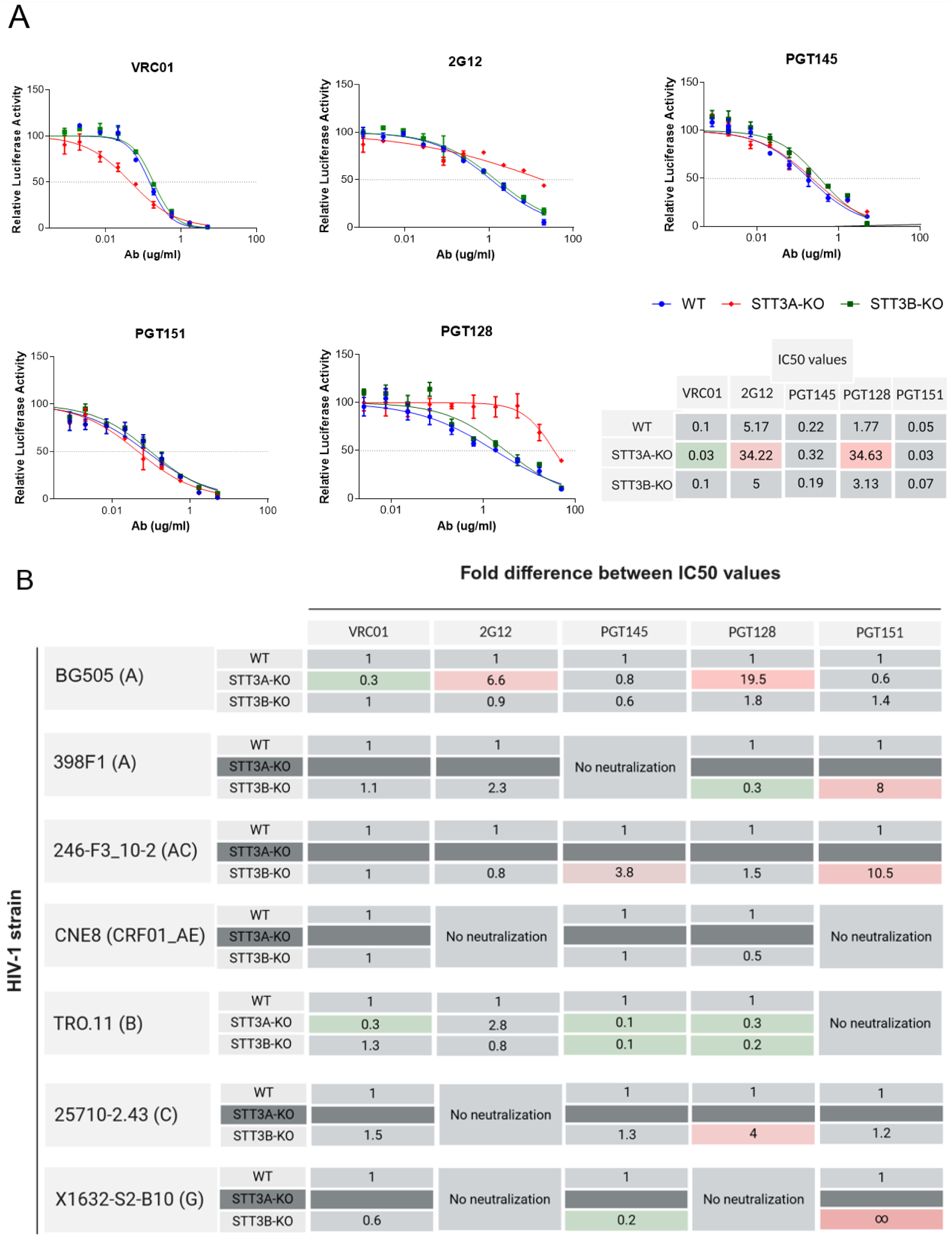
Effect of STT3A and STT3B knockouts on neutralization sensitivity against a panel of bNAbs. **(A)** Neutralization curves and IC_50_ values for the BG505 virus produced from an intact molecular clone and measured in TZM-bl reporter assay. IC_50_ values were calculated by fitting neutralization curves to a non-linear regression model and determining the antibody concentration required for 50% inhibition of infection. Changes in neutralization sensitivity are indicated by color coding: green represents a lower IC_50_ (increased sensitivity to neutralization), pink represents a higher IC_50_ (increased resistance to neutralization). Bars represent the mean ± standard deviation from at least three independent experiments. **(B)** The effect of STT3A/3B-KO on IC_50_ values for pseudoviruses from different clades is here reported as fold change in IC_50_ compared to pseudoviruses produced in wild type (WT) cells. The color coding used for the table in panel (A) was also applied to panel (B). Dark grey indicates that the virus produced in the respective cell lines was not sufficiently infectious to perform neutralization assays. For HIV-1 strains, the clade they belong to is indicated in parenthesis.

Subsequently, the neutralization sensitivity of our HIV-1 strains panel produced as pseudoviruses in the same cell lines was evaluated. Some strains produced in WT cells were resistant to specific bNAbs diminishing the value of the comparison between producer cells (Figure 3B). Only the TRO11 strain (clade B) pseudovirus was sufficiently infective when produced in STT3A-KO cells to proceed with neutralization experiments. TRO11 pseudovirus was more sensitive to VRC01 compared to its WT counterpart, similar to what was observed for BG505 virus. In contrast to BG505 virus, TRO11 produced in STT3A-KO cells was more sensitive to PGT128 neutralization, suggesting that glycans at positions N295, N301 and N332 that are part of the epitope it targets were attached. We speculate that neighboring glycans may be missing, providing a potential explanation for the enhanced neutralization observed.

STT3B-KO had no effect on the sensitivity of viruses to VRC01 or 2G12. When studying the STT3B-KO impact on PGT128 neutralization, while in some instances increased neutralization sensitivity was observed (i.e. 398F1, TRO.11), it also rendered the 25710-2.43 strain more PGT128 neutralization resistant. These alterations in sensitivity may be attributed to differential glycan occupancy at various sites within and surrounding the epitope. Furthermore, we observed that for several isolates the PGT151 targeted epitope was affected by STT3B-KO production, making 398F1 (clade A), 246-F3_C10_2 (clade AC), and X1632-S2-B10 (clade G) strains more resistant. We attribute this resistance pattern to the potential absence of interface glycans N611 and/or N637 that are required for efficient PGT151 binding. The effects of STT3A/3B-KO on PGT145 neutralization was also variable: while STT3B-KO increased sensitivity of TRO-11 and X1632-S2-B10, it rendered the 246-F3_C10_2 strain more resistant. These observations reveal that STT3A and STT3B ablation have highly variable effects on the neutralization sensitivity of different HIV-1 strains to bNAbs. Considering the function of the targeted enzymes knocked out and the bNAbs glycan specificities, the observed effects probably depend on whether glycans within the epitopes or around the epitopes targeted are absent. The former would be expected to the lead to resistance because critical epitope component are absent, while the latter could lead to enhanced access and therefore enhanced sensitivity.

### Knocking out STT3A or STT3B impacts trimerization

The occupancy of PNGS on recombinant HIV-1 Env trimers have been extensively studied, as has the composition of each glycan (48–55). We produced stabilized (SOSIP.v4.1; (56)) Env trimers of the BG505 strain used above, in WT, STT3A-KO and STT3B-KO HEK293T cell lines to study occupancy on individual PNGS found in BG505 Env. Two different purification methods were used. First, we used PGT145-antibody affinity chromatography, which is selective for well-folded native-like trimers (57). However, since PGT145 is dependent on glycans, we surmised that its use could bias the glycoforms selected after purification. The second method we used was therefore independent of glycosylation and involved Ni-NTA affinity chromatography followed by size exclusion chromatography (SEC).

Initial analysis of Ni-NTA-purified material using native gels demonstrated the presence of a heterogeneous mixture comprising trimers, dimers, and monomers, whereas PGT145 immuno-affinity purified material consisted exclusively of trimers (Figure 4A). This was expected, as tag-based purification methods capture all conformational variants of the protein. When subjected to SEC, a notable increase in the monomeric population was observed for proteins produced in STT3A-KO and STT3B-KO cell lines. This observation suggested that the absence of glycans at some sites may compromise the appropriate trimerization of Env protein. After SEC, the trimer peak was collected, and protein concentrations were measured for all three samples. The trimer yield from STT3A-KO cells was notably lower compared to both STT3B-KO and WT cells (Figure 4B).

**Figure 4.**
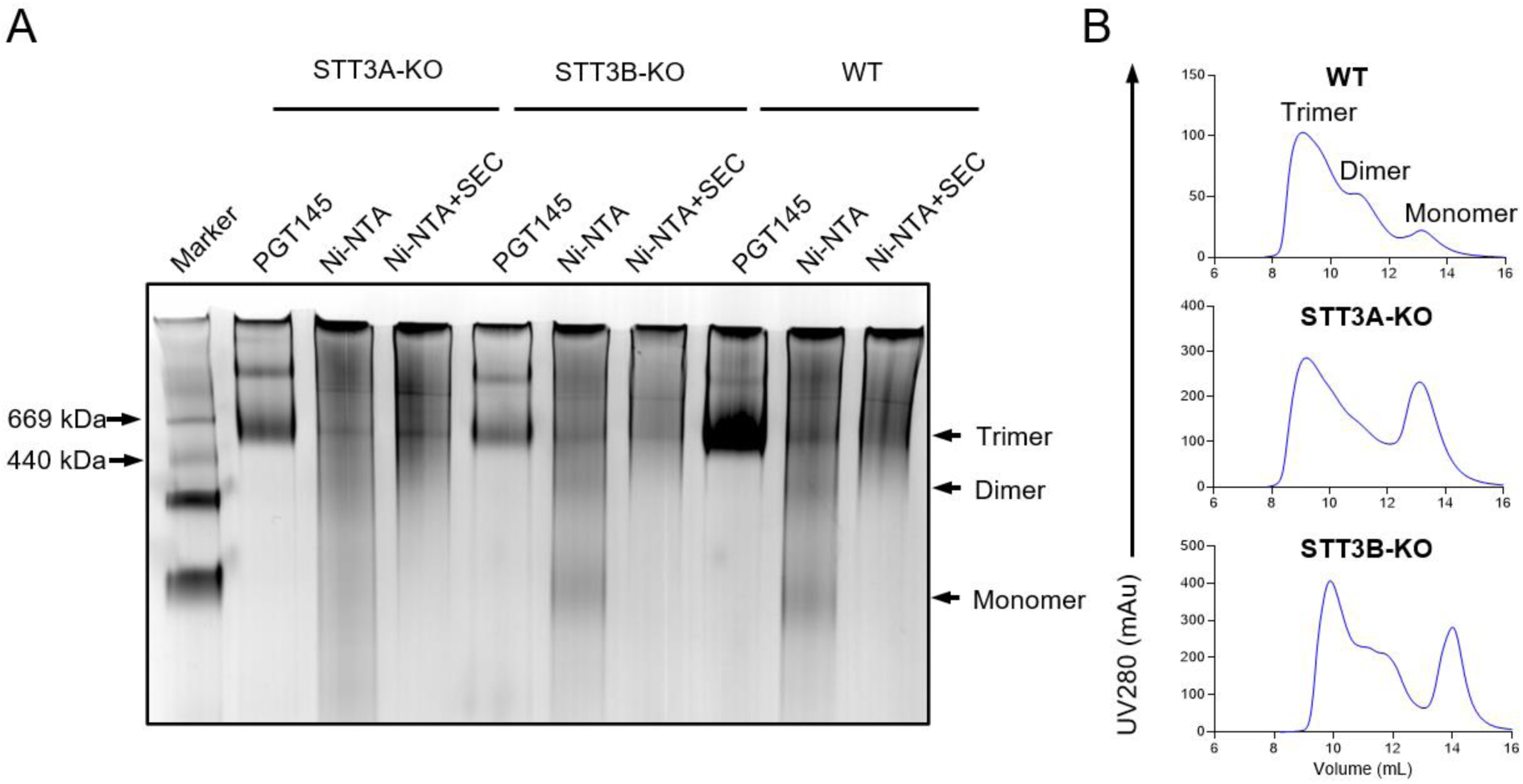
In vitro characterization of recombinant Env trimers produced in WT, STT3A-KO and STT3B-KO HEK293T cell lines. **(A)** Blue native-polyacrylamide gel electrophoresis analysis of Env proteins produced and isolated from different cell lines, stained by Coomassie blue. The PGT145- and Ni-NTA-purified (before and after SEC) Env proteins are shown. The molecular weight of two of the marker bands are indicated on the left end side of the gel picture (thyroglobulin and ferritin) and the expected positions for trimer, dimer, and monomer populations are indicated on the right end side of the picture. **(B)** SEC profiles of Ni-NTA purified Env proteins expressed in three different cell lines. A Superdex 200 10/300 GL column was used. The trimer, dimer, and monomer peaks are indicated. SEC, size exclusion chromatography.

### Knocking out of STT3A and STT3B affects binding of specific bNAbs

To further assess the impact of antigenicity of our various protein productions, we used biolayer interferometry (BLI) to investigate their binding properties against the panel of bNAbs described in the above sections. For the BLI assays, the proteins purified with PGT145 immuno-affinity chromatography were used, we note that this purification method is selective for well-folded native-like trimers, and deselects non trimeric and otherwise misfolded proteins species. High binding to quaternary- and glycan-dependent bNAb PGT151 suggested that some of the pre-fusion native-like features of the proteins produced from the three cells lines were preserved (Figure 5). Our data clearly showed PGT145 binding was found to be variable, which could be attributed to differences in the conformation of the apex, but likely also differences in occupancy of apex PNGS. BG505 SOSIP.v4.1 trimers produced in the absence of either STT3A or B did not display significant differences in binding to 2G12 and PGT128, irrespective of whether STT3A or STT3B was absent during production. In contrast, the binding of VRC01 was greater towards trimers produced in both STT3A-KO and STT3B-KO cells, in particular, this increase was highest for the STT3A -/- which is consistent with the enhanced neutralization sensitivity observed (Figure 3).

**Figure 5.**
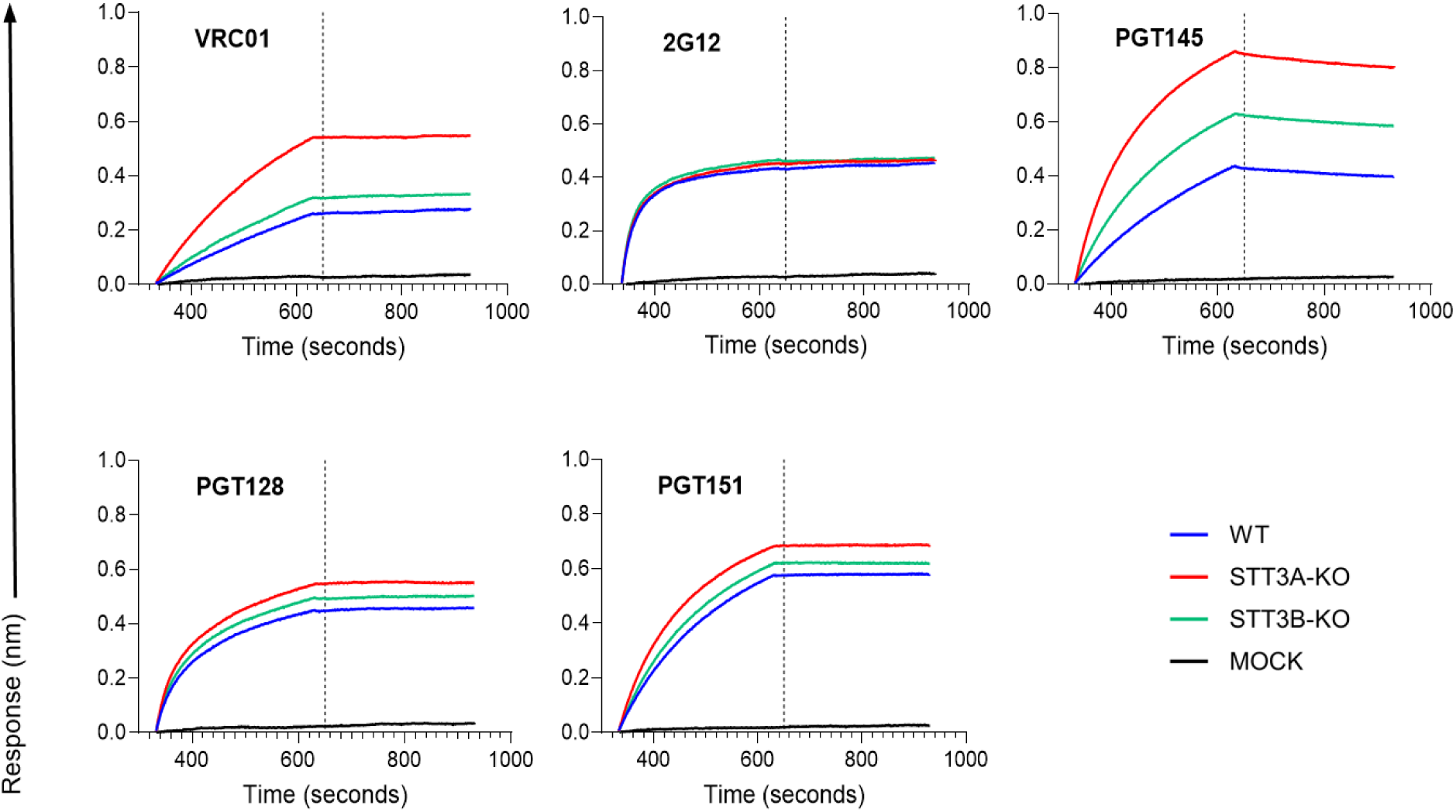
Binding of a panel of mAbs to Env proteins produced in different cell lines. BG505 SOSIP.v4.1 proteins were produced in wild type (WT), STT3A-KO or STT3B-KO HEK293T cell lines and purified through immune-affinity chromatography using PGT145. Biolayer interferometry (BLI) measurement were performed on the PGT145-purified trimers. The proteins were tested against five bNAbs (VRC01, 2G12, PGT145, PGT128, PGT151).

### Knocking out of STT3A and STT3B leads to under-occupancy of specific PNGS

Liquid chromatography-electrospray ionization (LC-ESI) mass spectrometry (MS) with an Orbitrap Fusion mass spectrometer was used to determine PNGS occupancy on the Env trimers (8). PNGS occupancy is expressed as the percentage of the total peptide that is modified by a glycan. Comparing the WT PGT145- and Ni-NTA/SEC-purified proteins showed similar PNGS occupancy patterns with a few minor exceptions where PGT145-purification appeared to have selected for more occupied PNGS (e.g. N190, N611 and N618) (Fig. 6A). Eliminating STT3A or STT3B resulted in less PNGS occupancy across the Env trimer. However, most PNGS remained highly occupied. This observation could stem from redundancy between OST-A and OST-B, leading to one isoform compensating for the absence of the other. The impact of STT3A/B deletion is likely more significant than what is observed from material purified post-production, as highly unoccupied material is prone to aggregation and will likely be degraded inside the cell.

**Figure 6.**
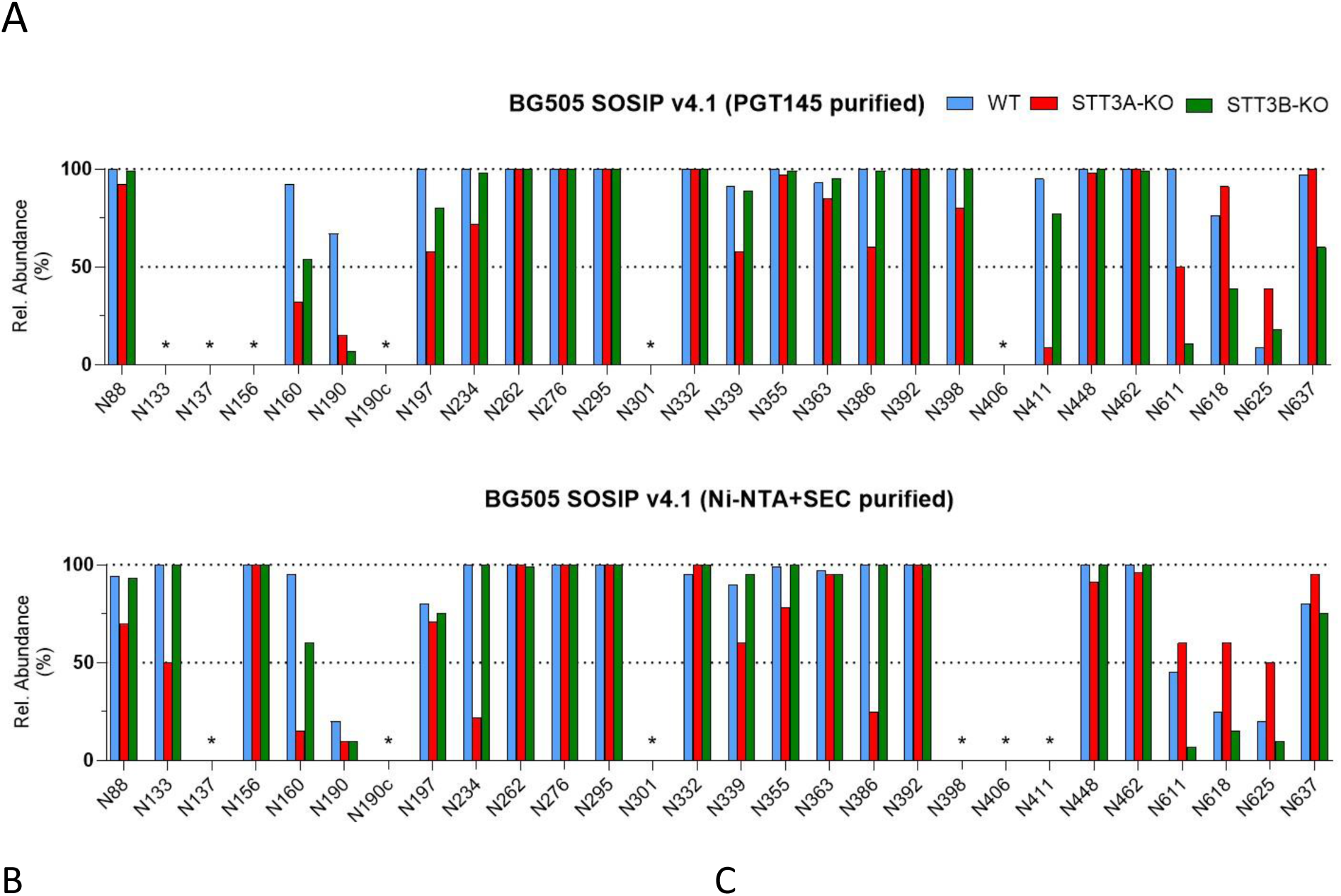
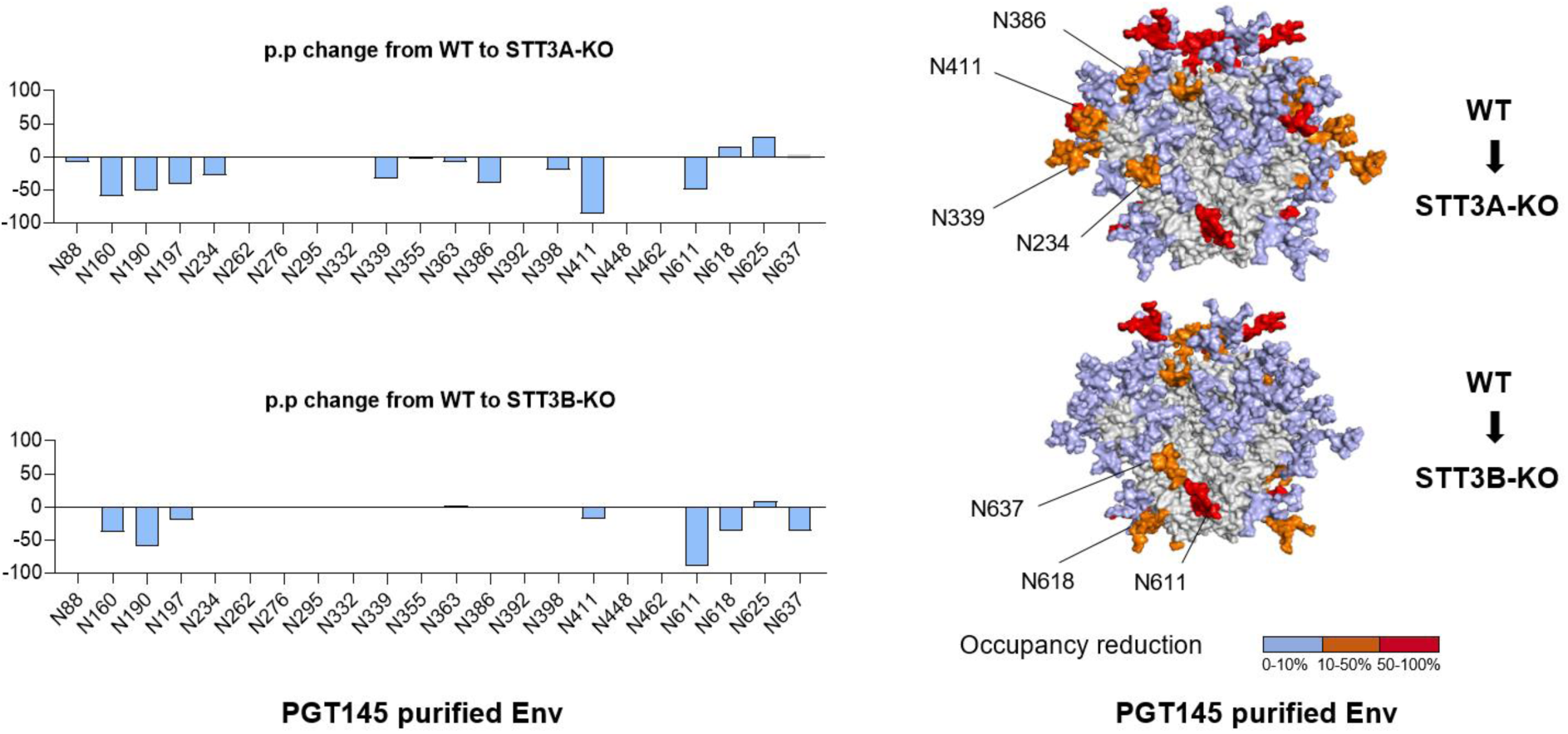
Glycan occupancy is decreased in STT3A-KO and STT3B-KO-produced BG505 soluble Env. **(A)** Quantification of site-specific occupancy for the 28 PNGS on Env trimers produced in different cell lines and analyzed by LC-ESI MS. The analysis included proteins purified via PGT145 immuno-affinity chromatography and those purified by Ni-NTA/SEC. Results are depicted as the mean of two independent biological replicates for each protein. The data displayed represents the PNGS occupancy expressed as the percentage of glycosylated peptide. ‘*’ indicates sites for which data could not be determined in at least one protein variant. **(B)** The presented data represents the arithmetic difference between the glycan occupancy of the STT3A- or STT3B-KO cell lines produced proteins minus the WT glycan occupancy, representing a percentage point change (p.p.). A negative p.p. change represents a lower occupancy of the KO variant compared to the WT. Only glycosylation sites for which data could be obtained for both WT and KO versions are included. **(C)** The structural models of the WT and KO Env trimers showing the glycan sites that were significantly impacted by STT3A and STT3B knockouts. Glycans are modeled onto each PNGS onto a previously generated BG505 structure (48). Colors indicate the degree of occupancy reduction from WT to STT3A/3B-KO: 0-10% (light blue), 10%–50% (orange), and 50%–100% (red).

When studying PGT145-purified trimers, STT3A-KO led to reduced occupancy (>20% reduction compared to WT) at PNGS at N160, N190, N197, N234, N339, N386, N411, and N611, while STT3B-KO led to less occupancy at N160, N190, N197, N611, N618, and N637. A few general observations can be made. First, there is substantial overlap in the sites that are underoccupied in both cell lines, including sites that PNGS closely spaced. For example N160, which is in close proximity with N156, and N190, which is in close proximity with N190c have reduced occupancy in both cell lines. N137, which is proximal to N133, also shows poor occupancy however the N137 containing glycopeptide was not resolved for the WT control. It is known that closely spaced PNGS are prone to under-occupancy (20) and our observations related to N160 and N190 suggest that this is exacerbated when STT3A or STT3B is absent. A decrease in occupancy at N197 was also observed in both KO cell lines (42% and 20% reductions for STT3A and STT3B-KO relative to WT, respectively), but the magnitude of this reduction was notably lower compared to that seen at N160 (60% and 38% reductions for STT3A and STT3B-KO relative to WT, respectively) and N190 (52% and 60% reductions for STT3A and STT3B-KO relative to WT, respectively) (Figure. 6A).

Subsequently, we studied whether particular glycosylation sites were dependent on either STT3A or STT3B. The glycan occupancy analysis showed that deletion of STT3A impacted glycosylation occupancy more globally than STT3B removal, even at sites such as N234, N339, N386, and N411 which are usually fully occupied (8, 20). These sites located on gp120 subunits can be considered STT3A-dependent, as a similar reduction in occupancy was not observed in proteins produced in STT3B-KO cells. STT3B-KO largely impacted the occupancy of C-terminal glycan sites N611, N618, and N637, found in the gp41 ectodomain of Env, and a reduction at these sites was not observed in the STT3A-KO version. These findings suggest that glycosylation at N611, N618 and N637 is preferentially mediated by STT3B, in line with literature demonstrating that STT3B is mostly responsible for adding glycans near the C-termini of glycoproteins (11, 16). This phenomenon may be in part explained by C-terminal sequons appearing late during translation, therefore residing for a shorter time in the translocation channel and limiting exposure to STT3A. The site-specific occupancy data for the PGT145-purified trimers were subsequently plotted as percentage point changes from WT to KO cells to highlight the differences in glycan occupancy at specific sites for glycan sites where occupancy could be determined experimentally (Figure 6B). To contextualize the changes in occupancy, we modeled glycans onto a previously generated WT structure (48) (Figure 6C).

## DISCUSSION

In this study, we investigated the role of STT3A and STT3B catalytic subunits on HIV-1 Env glycosylation in the context of Env-based vaccines. The OST-A and OST-B enzyme complexes, which contain STT3A and STT3B, respectively, add glycans to polypeptides cotranslationally or posttranslationally, respectively (13). It is now well known and studied that Env glycans hold a critical role in bNAb development during infection in people with HIV and that they have become a major focus of the HIV-1 vaccine design through their removal or addition to fill in glycan holes and/or create artificial glycan holes present on recombinant Env trimers used as immunogens to shape B cell responses towards neutralizing response (31, 37, 58, 59). However, the task is complex and controlling glycan occupancy often requires strain-specific engineering. Thus, developing new strategies may be needed to control glycan occupancy of Env-based vaccines. Defining the roles of N-linked glycosylation by STT3A and STT3B is beneficial in understanding the factors that result in the inefficient glycosylation of recombinant Env proteins. While the simultaneous knockout of STT3A and STT3B was not feasible because of cell death, the individual STT3A-KO or STT3B-KO cells helped to assess the role of each isoform in glycosylation. Overall, the current study demonstrated that the viral Env glycosylation is mostly controlled by and dependent on OST-A, and less so by OST-B. Although STT3A is primarily responsible for glycosylation of the viral Env, STT3B plays a comparatively greater role in the glycosylation of soluble Env immunogens. Furthermore, we found that the impact of STT3A/3B-KO on bNAb neutralization varied widely across different HIV-1 strains derived from different clades. Additionally, site-specific glycan analysis revealed that STT3A-KO results in a more global reduction in glycan occupancy, whereas STT3B plays a more specific role in the glycosylation of sites located closer to the C-terminus consistent with studies on cellular glycoproteins (16).

Our data on viral production and infectivity support that the OST-A complex plays a more critical role in Env glycosylation than OST-B. In the context of full-length HIV-1 Env presented on virions, the transmembrane domain positions the gp41 glycans significantly upstream of the gp41 cytoplasmic tail C-terminus. As a result, these glycans are not located within the region most sensitive to the STT3B-specific glycosylation pathway, which preferentially acts on sites closer to the C-terminus (16). Thus, the spatial separation of gp41 glycan sites from the C-terminal region provides a plausible explanation for the minimal impact of STT3B KO on viral infectivity, as most of the gp41 glycosylation would be handled by STT3A. The strong dependence on STT3A and the lack of compensation from STT3B can also be explained by the membrane-bound nature of the viral Env glycoprotein and the lack of a free C-terminus in the lumen of the ER. Given that Env is a membrane-bound protein that folds cotranslationally, the rapid folding process and its association with the membrane may limit the ability of STT3B to access and glycosylate skipped sites. Additionally, the repositioning of acceptor sites away from the ER lumenal surface during folding may further hinder STT3B- mediated modification (17).

Viral production data indicated that STT3B-KO enhanced 398F1 and X1632-S2-B10 production dramatically. However, this effect was only partially reflected in infectivity with 398F1 showing enhanced and X1632-S2-B10 no change in infectivity. Despite high levels of p24, infectivity remained low for X1632-S2-B10 strain, suggesting that a substantial proportion of the produced virions were non-infectious. In the context of pseudoviruses, a high p24 signal may reflect non-functional particles, such as those with impaired Env incorporation or conformational defects (60, 61). In contrast to these two strains, the 25710- 2-43 strain infectivity improved despite no significant change in viral production. These strain-specific outcomes highlight the complex interplay between glycosylation, Env incorporation, and viral fitness.

Another important consideration is that neutralization assays provide critical insights into the antigenic landscape of functional HIV-1 Env trimers. However, they inform on functional Env only and offer no information about non-functional Env that may also be present on the viral surface. As a result, differences in glycosylation and/or epitope exposure that exist on non-functional Env remain undetected in standard neutralization readouts. Additional assays such as binding to virion-derived Env or site-specific glycopeptide analysis can provide complementary resolution (62–64).

We reported enhanced VRC01 binding to Env produced in the absence of STT3A and increased neutralization sensitivity of pseudovirus generated in the absence of STT3A. Both results are consistent with improved accessibility of the CD4bs. We hypothesize that this is due to reduced occupancy of PNGS proximal to the CD4bs, in particular the PNGS at N190, N197, N234 and N386. This is in lines with observations that the absence of these sites improves epitope accessibility and increased neutralization sensitivity (65–69). Site-specific glycan analysis of the STT3A-KO Env protein supported this hypothesis, showing notable underoccupancy at positions N197 and N386 in particular.

In contrast, ablating STT3A activity conferred increased resistance to glycan-dependent bNAbs 2G12 and PGT128 for the BG505 strain. Here, we hypothesize that this resistance may result from underoccupancy at glycan sites that are part of the outer domain intrinsic mannose patch (46, 70, 71). Consistent with this observation, site-specific glycan analysis of the soluble BG505 SOSIP.v4.1 purified with PGT145 revealed reduced occupancy at several relevant PNGS, including N339 and N386, which are known to contribute to the epitopes recognized by V3-glycan-targeting bNAbs. Altogether, these findings suggest that STT3A- mediated glycosylation is critical for either shielding critical epitopes from antibody recognition or maintaining sensitivity to certain glycan-dependent antibodies especially for BG505 Env protein and virus.

Site-specific glycan analysis of recombinant Env trimers showed that even when expressed in WT cells, purification strategy can significantly affect the glycan occupancy of soluble Env trimers. Notably, purification via PGT145 affinity chromatography led to >20% higher occupancy at glycan sites N190, N197, N611 and N618 compared to Ni-NTA/SEC purification when expressed in WT cells. As glycosylation is a critical determinant of Env antigenicity and immunogenicity, these differences underscore the importance of carefully selecting a purification method, particularly in the context of immunogen production.

Glycan analysis revealed that a large majority of the PNGS were still highly occupied from all producer cell lines tested. The ability to preserve glycan occupancy despite OST deficiencies highlights that the OST isoforms have redundant roles. This observation is, however, made for secreted purified glycoproteins, i.e. glycoproteins that have successfully completed the chaperone-assisted folding pathway. Our assays do not allow analysis of glycoproteins that did not fold properly because critical glycans were missing and were targeted for degradation (72, 73). Similarly, our viral production and infectivity analysis were performed on secreted viral particles. It is important to note that this may result in an overestimation of glycan occupancy as misfolded and/or under-glycosylated Env forms are less likely to be secreted or incorporated into virions.

Our results show that maximal glycosylation of Env involves the cooperation of both OST isoforms and this is consistent with observations with other glycoproteins (13). Viral Env proteins are crucial for inducing neutralizing antibodies due to their critical involvement in viral entry and their potential as immunogen candidates (74–77). Therefore, fundamental understanding of the role of OST complexes in viral envelope glycosylation could potentially lead to the development of new methods to mimic a fully glycosylated viral envelope and improve antigen quality, stability and relevance. In HIV-1 vaccine research, the design of Env trimer immunogens has taken a central role with the aim at inducing broad and protective neutralizing antibody responses. To ensure that recombinant Env trimers mimic viral Env in terms of glycan occupancy, artificial glycan holes should be eliminated by increasing PNGS occupancy and membrane-bound immunogen approaches favored (8). These adjustments hold promise with regard to immunogen production batch to batch consistency as well as controlling glycan hole as needed to open and close areas on Env trimers to either enhance immune-focusing efforts or to further guide immune responses towards breadth mimicking antibody-virus co-evolution in people living with HIV that developed neutralization breadth and/or bNAbs linked to glycan shield evolutionary events. Revealing how OST-A and OST-B complexes contribute to the Env glycosylation offer valuable insights for immunogen design and will likely facilitate the development of methods, producer cells and protein engineering to control glycan occupancy.

## MATERIALS AND METHODS

### Cell culture

Recombinant Env proteins and the infectious virus stocks were prepared by transfecting WT HEK293T (American Type Culture Collection Cat. #11268) as well as HEK293T STT3A-KO, and STT3B-KO cell lines kindly provided by Dr. Reid Gilmore and Dr. Natalia Cherepanova. Cells were cultured under sterile condition and kept at +37°C, 5% CO_2_, in a humidified incubator. Cell lines were maintained in culture using DMEM plus glutamate (GIBCO) and 10% fetal calf serum (FCS) supplemented with antibiotics (i.e. penicillin and streptomycin, 100 U/ml each)(complete medium) and 0.05% w/v of Trypsin/EDTA solution was used to detach and passage the cells.

### BG505 intact virus and pseudovirus production

For the generation of intact virus stocks, WT HEK293T (2 × 10⁵), STT3A-KO (2.5 × 10⁵), and STT3B-KO (2 × 10⁵) cells were seeded in 3 mL/well of complete medium in 6-well tissue culture plates (Corning) to reach ∼85–90% confluency at the time of transfection. Cells were transfected with 5 µg of previously described BG505 strain plasmid genomic DNA (20). To produce Env-pseudotyped viruses, WT and KO HEK293T cells were seeded at the same densities as for BG505 intact virus production. Transfections were carried out using 1.6 µg of a full-length Env expression plasmid and 2.4 µg of the Env-deficient HIV-1 backbone plasmid pSG5ΔEnv, as previously described (78). For both productions, plasmid DNA was diluted in 250 µL Opti-MEM (GIBCO) and mixed with 10 µL of Lipofectamine 2000 (Invitrogen) that had been pre-diluted in 240 µL Opti-MEM. After a 20-minute incubation at room temperature, transfection mix solutions were added to the cells. Supernatants containing virus were harvested 48 hours post-transfection and filtered through a 0.45 µm membrane for downstream use.

### p24 Capsid (CA-p24) ELISA

To quantify HIV-1 particles, production of the capsid protein p24 (CA-p24) was assessed using the HIV-1 Gag p24 DuoSet ELISA kit (Bio-Techne, R&D Systems) following a custom in-house protocol (41). High binding half-area white 96-well plates (Greiner Bio-One) were coated with mouse anti-HIV-1 Gag CA-p24 capture antibody and incubated overnight at room temperature. The following day, plates were washed three times with wash buffer (1X PBS + 0.05% Tween 20) then blocked with 1% BSA in 1X PBS, 0.2% Triton X-100. Diluted samples, CA-p24 standard proteins, and controls (PBS) were added to the wells and incubated for 2 hours at room temperature with continuous shaking. After incubation, the wells were washed again using washing buffer. A 1:80 dilution of Streptavidin-HRP (Bio-Techne, R&D Systems) was prepared using Reagent Diluent (Bio-Techne, R&D Systems) and added to the wells, followed by a 20-minute incubation at room temperature. The plates were then washed again and tapped dry. Finally, a 1:10 dilution of LumiPhos A+B substrate was prepared using Milli-Q water and added to the wells for 2 minutes to allow signal development Luminescence was measured immediately using a GloMax® plate reader (Promega). Standard curves were generated, and data analysis was conducted to determine CA-p24 concentrations. The various productions of HIV-1 pseudovirus isolates and BG505 virus were measured in duplicate and in two independent ELISAs.

### Infectivity assay

TZM-bl cells (79) were seeded at a density of 1.7 × 10⁴ cells per well in 96-well plates one day before infection. Cells were cultured in complete medium and passaged as describe above. On the following day, the harvested intact and pseudoviruses were titrated using TZM-bl cells and in quadruplicate as previously described (79). The TCID_50_ values were determined for each virus using Graphpad Prism v10.

### Antibody production

Recombinant antibodies were produced in HEK293F suspension cells as previously described (44). Briefly, heavy and light chain plasmid DNAs (156 μg each) were filtered and combined with polyethylenimine (PEI) MAX (Polysciences) at a 1:3 DNA:PEI ratio in Opti-MEM reduced serum medium (Gibco). The mixture was incubated for 30 minutes at room temperature before being added to HEK293F cultures. After a 6–7 day incubation, supernatant were collected, centrifuged, and filtered through 0.2 μm filters. Protein G coupled Agarose beads (Pierce, 20397) were added to the filtered supernatants, incubated overnight at +4 °C with gentle rotation, the resin transferred to a centrifuge column (Pierce), washed twice with PBS (pH 7.2), and antibodies eluted using 0.1 M glycine (pH 2.5). The eluate was immediately neutralized using 1 M Tris (pH 8.6) as collection buffer in a 9:1 ratio. The eluate were concentrated and exchanged into 1X PBS using a 100 kDa molecular weight cut-off Vivaspins (Sartorius) and then passed through a 0.22 μm filter (Costar, 98231-UT-1). Final protein concentrations were determined by UV280 absorbance using the standard IgG extinction coefficient on a Nanodrop 2000 instrument (Thermo Scientific).

### Neutralization assay

TZM-bl cells were seeded at a density of 1.7 × 10⁴ cells per well in 96-well plates. The following day, the viral input equal to TCID_50_ for each virus was incubated for 60 min at room temperature using 3-fold serial dilutions of each bNAb tested. This mixture supplemented with 40 µg/mL DEAE-dextran (40 μg/mL) and saquinavir (400 nM) at a 1:1 ratio were added onto TZM-bl cells to a final volume of 200 µL/well. Three days post-infection, cells were washed with 1X PBS and lysed using lysis buffer (25 mM Glycylglycine (Gly-Gly), 15 mM MgSO4, 4 mM EGTA tetrasodium, 10% Triton X-100, pH 7.8). Bright-Glo kit (Promega, Madison, WI) was used to measure Luciferase activity. All infections were carried out in quadruplicate to ensure reproducibility. Background luminescence was subtracted using uninfected control wells. For normalization, the infectivity of each Env mutant in the absence of antibody was defined as 100%. Dose–response curves were generated using nonlinear regression, and the half-maximal inhibitory concentration (IC_50_) values were determined by fitting a sigmoidal curve in GraphPad Prism v10.

### Blue Native-PAGE

To assess the correct folding into trimers of the produced proteins, these were subjected to a Blue Native-Polyacrylamide gel electrophoresis and visualized using colloidal blue stain (Life Technologies). Typically, 3 μg of purified protein was prepared in 4× MOPS buffer (200 mM MOPS, 200 mM Tris, pH 7.7) and resolved on a NuPAGE 4–12% Bis-Tris gel (Novex). Electrophoresis was carried out for up to 60 minutes at 200 V using Invitrogen cathode (NB2001) and anode (NB2002) buffers. Gels were then stained with colloidal blue stain following the manufacturer’s instructions. Once destained with MiliQ water, the gels were imaged using a gel imaging system (Bio-Rad).

### Biolayer Interferometry (BLI)

Antibody binding to PGT145-purified Env trimers produced in WT, STT3A-KO, and STT3B-KO HEK293T cells was analyzed using a ForteBio Octet K2 instrument, as previously described (80). All measurements were performed at +30°C with shaking at 1000 rpm. Antibodies and purified proteins were diluted in running buffer (1X PBS containing 0.1% BSA and 0.02% Tween-20) to a final volume of 300 μL/well. Protein A biosensors (ForteBio) were loaded with antibodies at a concentration of 2.0 μg/ml until a loading threshold of 0.5 nm was reached. Trimer proteins were prepared at 600 nM with association and dissociation phases monitored for 300 seconds each. Background binding was determined using sensors loaded with trimer plunged into running buffer in the absence of antibody.

### Env SOSIP trimers

The BG505 SOSIP.v4.1-derived trimers have been extensively described elsewhere, as have the methods to produce and purify them (56). Briefly, Env trimers were produced in WT HEK293T (15 × 10⁶), STT3A-KO (18.6 × 10⁶), and STT3B-KO (15 × 10⁶) cells seeded into multilayer T175 flasks (Thermo Scientific™). Each flask was transfected with 180 μg of plasmid encoding the HIV-1 Env protein of interest and 50 μg of furin expression plasmid using PEI MAX. Following 72h transient expression, supernatants were harvested and Env trimers purified either by PGT145-immunoaffinity chromatography (56) or Ni-NTA affinity chromatography followed by size exclusion chromatography (SEC) (Bio-Rad) to check for quality and isolate trimer peaks.

### Site-specific glycan analysis using mass spectrometry

For protein samples to be analyzed by LC-MS, three separate 50 μg aliquots were denatured for 1 h in 50 mM Tris/HCl, pH 8.0 containing 6 M of urea and 5 mM of dithiothreitol (DTT). Next, the proteins were reduced and alkylated by adding 20 mM iodoacetamide (IAA) and incubated for 1 h in the dark, followed by incubation with DTT to get rid of any residual IAA. The alkylated proteins were buffer-exchanged into 50 mM Tris/HCl, pH 8.0 using Vivaspin columns (3 kDa), and digested separately overnight using trypsin, chymotrypsin (Mass Spectrometry Grade, Promega) or alpha lytic protease (Sigma-Aldrich) at a ratio of 1:30 (w/w). The peptides were dried and extracted using Oasis HLB 96-well plates (Waters). The peptides were dried again, re-suspended in 0.1% formic acid and analyzed by nanoLC-ESI MS with an Easy-nLC 1200 (Thermo Fisher Scientific) system coupled to an Orbitrap Fusion mass spectrometer (Thermo Fisher Scientific), using stepped higher energy collision-induced dissociation (HCD) fragmentation (15, 25, 45%). Peptides were separated using an EasySpray PepMap RSLC C18 column (75 µm × 75 cm). A trapping column (PepMap 100 C18 3 μm (particle size), 75 μm × 2 cm) was used in line with the LC prior to separation with the analytical column. The LC conditions were as follows: 275 min linear gradient consisting of 0– 32% acetonitrile in 0.1% formic acid over 240 min followed by 35 min of 80% acetonitrile in 0.1% formic acid. The flow rate was set to 200 nL/min. The spray voltage was set to 2.7 kV and the temperature of the heated capillary was set to 40 °C. The ion transfer tube temperature was set to 275 °C. The scan range was 400−1600 m/z. The HCD collision energy was set to 50%. Precursor and fragment detection were performed using Orbitrap at a resolution MS1 = 100,000. MS2 = 30,000. The AGC target for MS1 = 45 and MS2 = 54 and injection time: MS1 = 50 ms, MS2 = 54 ms.

### Data processing of LC-MS data

Glycopeptide fragmentation data were extracted from the raw file using Byos v3.5 (Protein Metrics Inc.). The following parameters were used for data searches in Byonic: The precursor mass tolerance was set at 4 ppm and 10 ppm for fragments. Peptide modifications included in the search include: Cys carbamidomethyl, Met oxidation, Glu pyroGlu, Gln pyroGln and N deamidation. For each protease digest, a separate search node was used with digestion parameters appropriate for each protease (Trypsin RK, Chymotrypsin YFW and ALP TASV) using a semi-specific search with 2 missed cleavages. A 1% false discovery rate (FDR) was applied. All three digests were combined into a single file for downstream analysis. All charge states for a single glycopeptide were summed. The glycopeptide fragmentation data were evaluated manually for each glycopeptide; the peptide was scored as true-positive when the correct b and y fragment ions were observed along with oxonium ions corresponding to the glycan identified. The relative amounts (determined by comparing the XIC of each glycopeptide, summing charge states) of each glycan at each site as well as the unoccupied proportion were determined by comparing the extracted chromatographic areas for different glycotypes with an identical peptide sequence.

## ACKNOWLEDGEMENTS

We thank Dr. Reid Gilmore and Dr. Natalia Cherepanova for providing HEK293T STT3A-KO and STT3B-KO cell lines. This work was supported by the U.S. National Institutes of Health Grant P01 AI110657 (to A.B.W. and R.W.S.); and by the Bill and Melinda Gates Foundation through the Collaboration for AIDS Vaccine Discovery (CAVD), grants INV-002022 and INV-063951 (to R.W.S.) and INV-070116 (to M.C.).

